# blupADC: An R package and shiny toolkit for comprehensive genetic data analysis in animal and plant breeding

**DOI:** 10.1101/2021.09.09.459557

**Authors:** Quanshun Mei, Chuanke Fu, Jieling Li, Shuhong Zhao, Tao Xiang

**Affiliations:** Key Laboratory of Agricultural Animal Genetics, Breeding and Reproduction of Ministry of Education & Key Laboratory of Swine Genetics and Breeding of Ministry of Agriculture, Huazhong Agricultural University, Wuhan 430070, China

## Abstract

**Summary:** Genetic analysis is a systematic and complex procedure in animal and plant breeding. With fast development of high-throughput genotyping techniques and algorithms, animal and plant breeding has entered into a genomic era. However, there is a lack of software, which can be used to process comprehensive genetic analyses, in the routine animal and plant breeding program. To make the whole genetic analysis in animal and plant breeding straightforward, we developed a powerful, robust and fast R package that includes genomic data format conversion, genomic data quality control and genotype imputation, breed composition analysis, pedigree tracing, analysis and visualization, pedigree-based and genomic-based relationship matrix construction, and genomic evaluation. In addition, to simplify the application of this package, we also developed a shiny toolkit for users.

**Availability and implementation:** blupADC is developed primarily in R with core functions written in C++. The development version is maintained at https://github.com/TXiang-lab/blupADC.

**Supplementary information:** [Supplementary data](https://github.com/TXiang-lab/blupADC/blob/master/Document_en.pdf) are available online

## 1 Introduction

A powerful genomic breeding software requires good organization of different genetic analyses and can be used for handling with different types of data in the whole process of breeding program. With a widely application of genomic selection (GS) in animal and plant breeding, the complications of involved genetic analyses keep increasing (Hickey, et al., 2017). Such situation is being further exacerbated with the increased demand of complex breeding strategy in combination with newly developed algorithms(Legarra, et al., 2015; Misztal, et al., 2015). Thus, to fulfill the high demand of handling with large-scale breeding datasets in high speed and optimized memory, some genomic evaluation software, such as two well-known software DMU (Madsen and Jensen 2013) and BLUPF90(Misztal, et al., 2002), were developed by compiled programming languages, including Fortran and C++.

On the other hand, the usage of genetic evaluation software is also a severe limitation for small-scale farms where an independent breeding program needs to be executed. Although there are some genetic evaluation software available in animal and plant breeding over 20 years, such as DMU and BLUPF90, users still must manually prepare all the datasets in a certain manner, do genomic quality control, genotype imputation, pedigree tracing, relationship matrix construction etc., and users need to be well trained before setting up a driver file for using these software, which is complicated and error-prone, even for experienced staffs.

R programming language (Team, 2018) is an open-source programming language that is widely used as a statistical software and data analysis tool. The R programming language has gained a lot of popularity recently because it is user friendly and it enjoys a vast Community Support.

Hence, we develop and present an R package and shiny toolkit, blupADC, aiming at assisting breeders to easily analyzing genomic data (including genomic data format conversion, data quality control and genotype imputation, breed composition analysis, pedigree tracing and visualization, relationship matrix construction etc.) before implementing genomic evaluation, and simplifying the procedure of running breeding program (including breeding model construction and automatically interface with breeding software DMU and BLUPF90) in animal and plant breeding systems.

## 2 Implementation

blupADC is implemented in R programming language (Team, 2018). To accelerate the computation speed, core functions in blupADC are written in C++ via Rcpp (Eddelbuettel, et al., 2011). Linear algebra calculations, such as matrix decomposition and matrix multiplications, are achieved by Armadillo C++ library via RcppArmadillo (Eddelbuettel and Sanderson, 2014). To handle with large-scale genomic data, multithreading is achieved by using OpenMP in C++ functions (Chandra, et al., 2001). In addition, as shown in **Figure 1**, to simplify the application of this package, we also developed a shiny toolkit for users(http://47.95.251.15:443/blupADC/).

**Figure 1.**
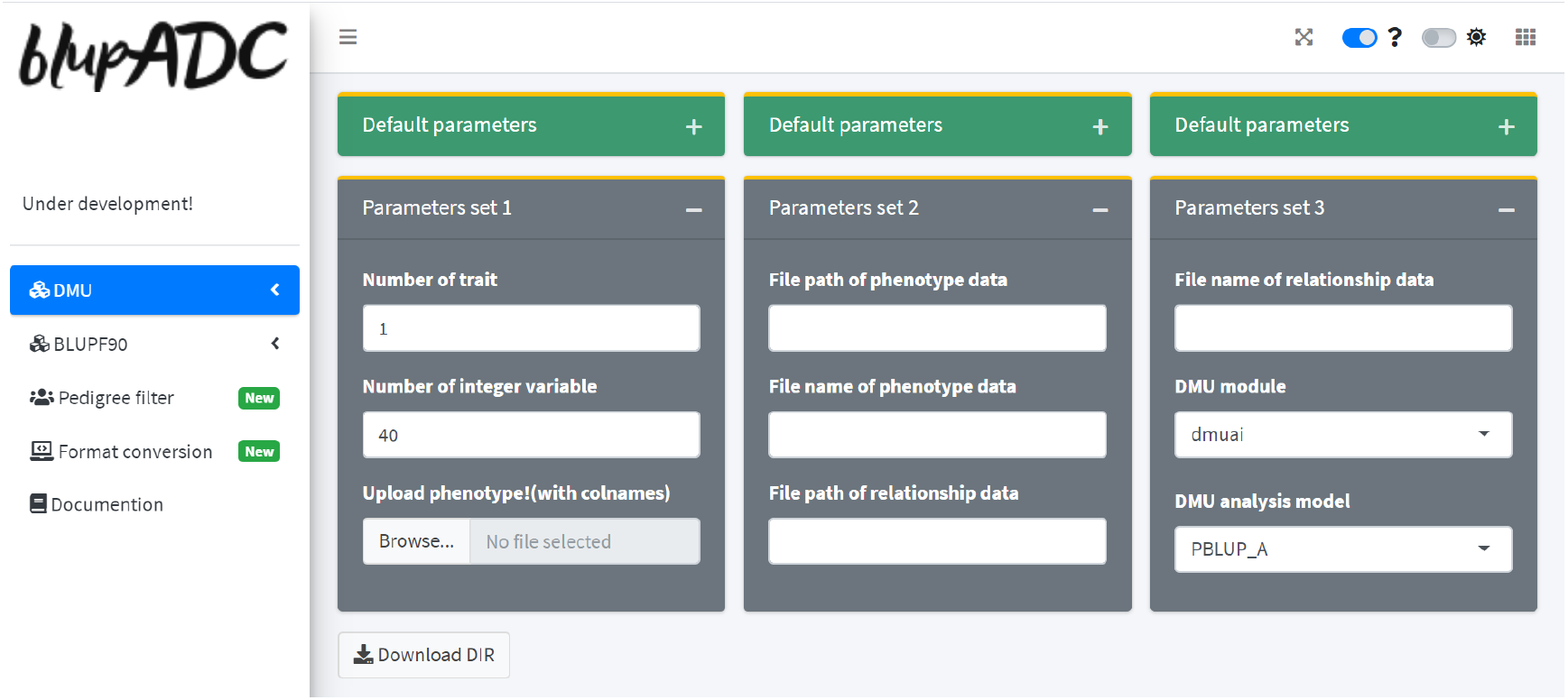
Shiny application of package blupADC.

Functions in blupADC can be grouped into 6 categories on the basis of its functionalities (**Figure 2**):

**Figure 2.**
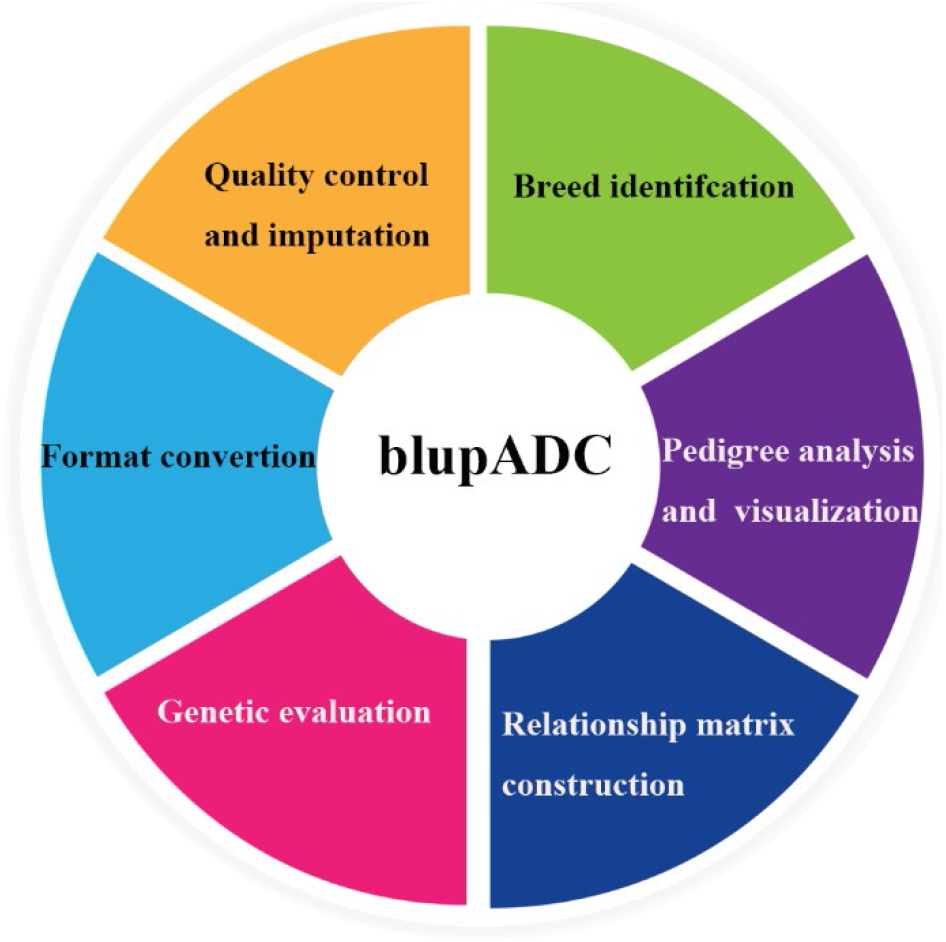
Overview of the blupADC package functionality.

- Genomic data format conversion
- Genomic data quality control and genotype imputation
- Breed composition analysis
- Pedigree tracing, analysis and visualization
- Relationship matrix construction
- Genetic evaluation

The Supplementary Materials contain detailed information on the usage of blupADC.

## 3 Features

### 3.1 Genomic data format conversion

blupADC supports multiple formats conversion for genomic data, including Hapmap(Gibbs, et al., 2003), Plink(Purcell, et al., 2007), BLUPF90(Misztal, et al., 2002), Numeric(VanRaden, 2008) and VCF(Siva, 2008) formats. These formats are commonly used for genomic data and they can also be used directly in most software of gnome-wide association studies and genomic selection, such as GCTA, EMMAX, and BLUPF90 (Kang, et al., 2010; Misztal, et al., 2002; Yang, et al., 2011).

### 3.2 Genomic data quality control and genotype imputation

Quality control and imputation for genomic data are the fundamental steps in genomic evaluation. Two robust software Plink(Purcell, et al., 2007), and Beagle(Browning, et al., 2018) have shown great advantages over many other software and been widely used in handling with genomic data and genotype imputation. However, to use Plink and Beagle appropriately, data format preparations and parameters setting are somehow tough. Thus, blupADC provides an R wrapper for interfacing with Plink and Beagle, which integrates the analyses of genomic data quality control and genotype imputation in a function, so that user can complete the relevant analyses easily.

### 3.3 Breed composition analysis

With the fast development of high-throughput genotyping techniques, breeders utilize SNP arrays to recognize population structure (Patterson, et al., 2006). Since principal component analysis (PCA) can help to investigate ethnic background of individuals (McVean, 2009), blupADC employs PCA to recognize population structure and uses k-means algorithm to cluster individuals with similar principal component values (Likas, et al., 2003). Individuals with error recording of breed can be automatically recognized and corrected.

### 3.4 Pedigree analysis and visualization

Pedigree information is the basis to explore the genetic effects in the animal and plant breeding. To study the target individuals, usually, a subset of the whole pedigree is of great interest for researchers. blupADC allows users to recode pedigree, correct common errors(offspring born before its parents, same offspring have different parents, and etc) in pedigree, and trace pedigree back for the target individuals for pre-set generations. In addition, blupADC provides an intuitive structure for pedigree by using the grammar of graphics implemented in R package ggplot2 (Wickham, 2011). As shown in **Figure 3**, each family structure can be identified and plotted automatically. Moreover, blupADC allows users to use abundant symbols to visualize the pedigree structure on the basis of R package ggimage (Yu, 2017).

**Figure 3.**
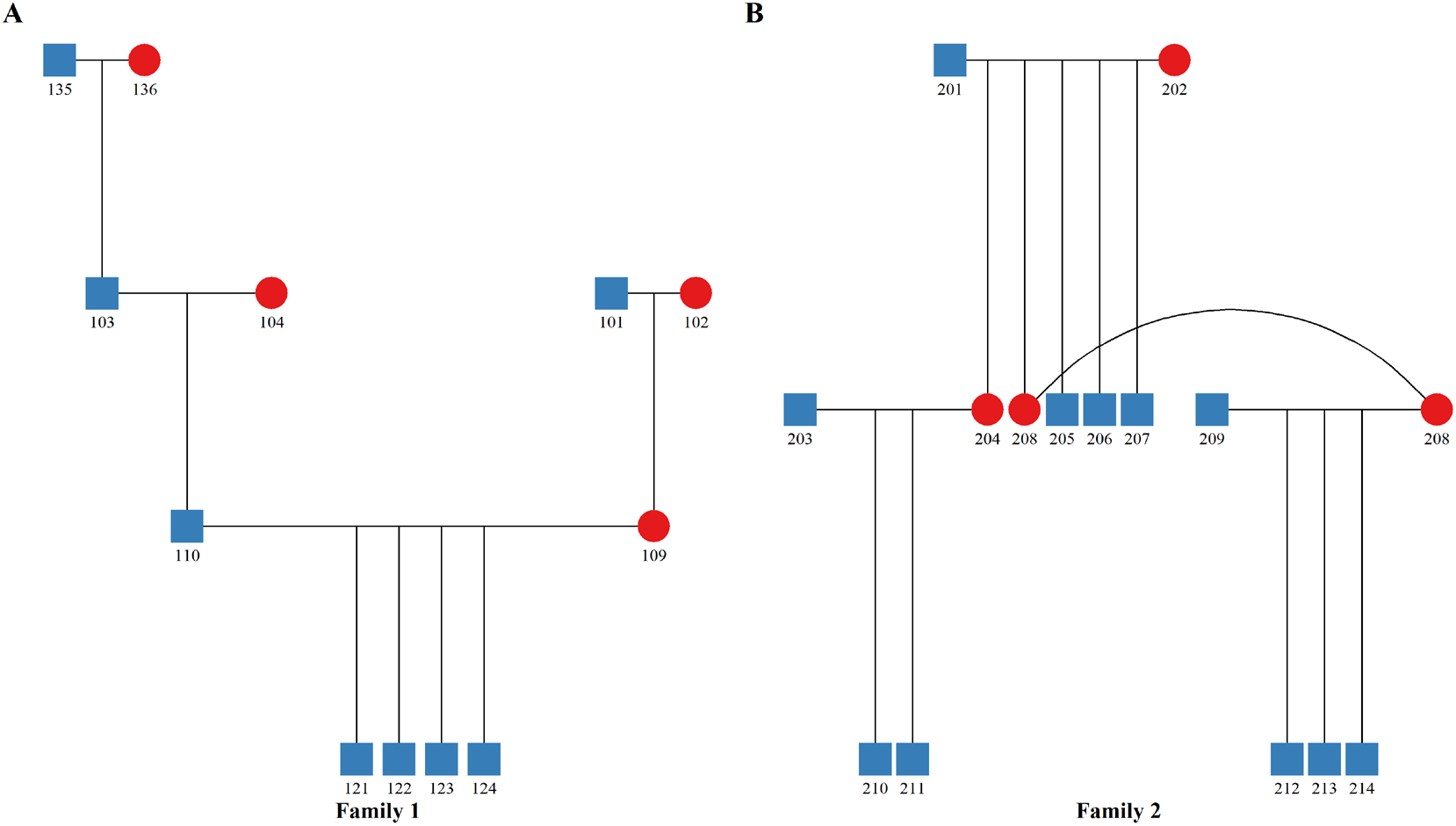
Pedigree structure of two families. A. Pedigree structure of family 1. B. Pedigree structure of family 2.

### 3.5 Relationship matrix construction

The classical algorithm best linear unbiased prediction (BLUP) was initially developed by Henderson in the 1950s, and it is widely used as the routine genetic evaluation methodology for animal and plant breeding (Gianola and Rosa, 2015; Hickey, et al., 2017). With the fast development of genotype sequencing technology and new algorithms, in the recent decades, there has been a growing interest in integrating genomic information with BLUP method. For example, single-step genomic best linear unbiased prediction (SSGBLUP) (Christensen and Lund, 2010; Legarra, et al., 2009), which has an ability to combine information of pedigree and genotypes to construct relationships for both genotyped and ungenotyped individuals, is commonly used in livestock breeding system. All of the genetic evaluation methods, which under the BLUP framework, need to provide a genetic relationship matrix to the used linear mixed model (LMM). Usually, these relationship matrices were constructed via using different information (e.g.: pedigree and genotypes) and algorithms. blupADC allows users to use different information to calculate (genomic) inbreeding coefficients and construct pedigree-based (Henderson, 1976), genomic-based (VanRaden, 2008) and the combined pedigree and genomic based (Legarra, et al., 2009) additive genetic relationship matrices for different genetic evaluation methods, including BLUP, Genomic BLUP and SSGBLUP. Furthermore, to capture dominance variance in the genetic evaluation, blupADC also allows user to construct pedigree-based dominance relationship matrix and genomic-based dominance relationship matrix. In terms of the pedigree-based dominance relationship matrix, blupADC provides two algorithms to construct it, including classical algorithm (Falconer and Mackay, 1996) and gene dropping algorithm (Mrode, 2014; Ovaskainen, et al., 2008). For the genomic-based dominance relationship matrix, blupADC allows users to construct two different types of matrices, which are also known as ‘classical’ (Su, et al., 2012) and ‘genotypic’ dominance relationship matrices (Vitezica, et al., 2013). In addition, blupADC provides options to construct two newly developed relationship matrices, including metafounder algorithm(Legarra, et al., 2015) and APY algorithm(Misztal, et al., 2015). The output of relationship matrix can be directly used by genetic evaluation software, including DMU (Madsen and Jensen, 2013) and BLUPF90(Misztal, et al., 2002).

### 3.6 Genetic and genomic evaluation

blupADC provides a R wrapper to interface with genetic evaluation software DMU and BLUPF90, which are popular and powerful in animal and plant genetic evaluation. blupADC automatically generated driver files for DMU and BLUPF90 on the basis of the entered parameters, and therefore users do not need to prepare the complicated and error-prone driver files by hand. In addition, after running genetic evaluation, blupADC provides tidy output files and makes the estimated genetic parameters visualized (**Figure 4**). Through using blupADC, user can easily accomplish genetic and genomic evaluations, including BLUP, Genomic BLUP, and SSGBLUP analyses with either single-trait or multiple-trait model. In addition to regular animal model, maternal effect model, permanent effect model, random regression effect model, and social genetic effect model can also be easy achieved. All the functions in DMU and BLUPF90 can be realized in a simple manner via using blupADC.

**Figure 4.**
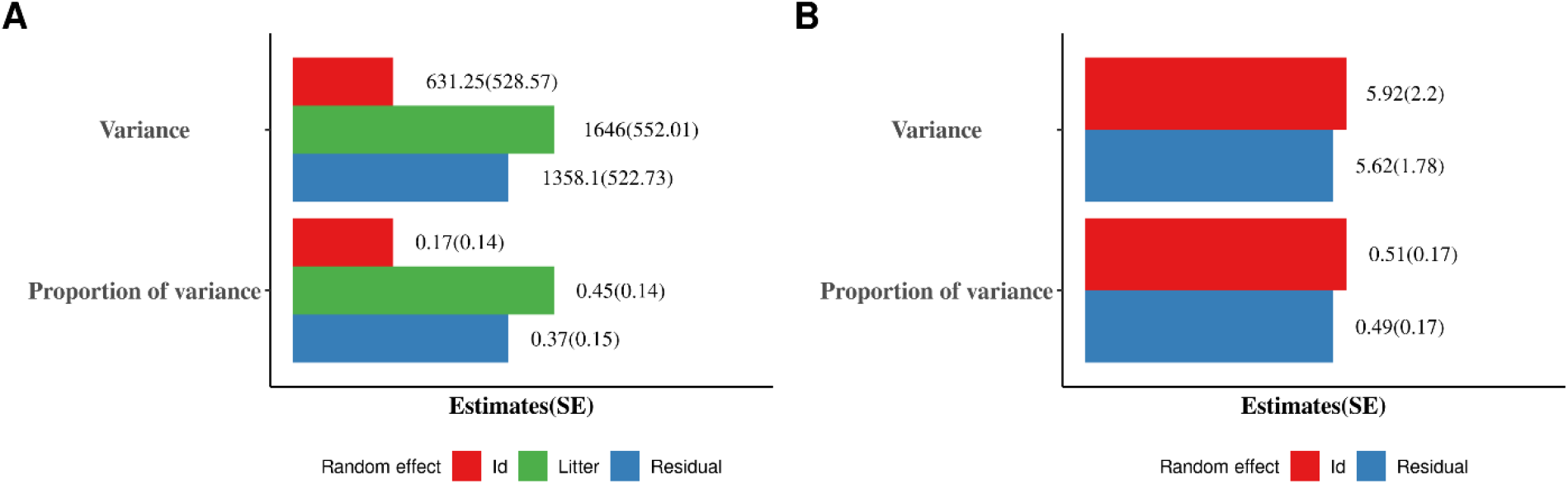
Variance components for two traits. A. Variance components of trait 1. B. Variance components of trait 2.

## Conclusions

blupADC is a robust and shiny toolkit for comprehensive genetic data analysis in animal and plant breeding. It allows users to perform all the essential steps in genomic breeding, including genotypic data format conversion, genotypic data quality control and genotype imputation, breed composition analysis, pedigree analysis and visualization, relationship matrix construction and execute genetic and genomic evaluation. blupADC simplify the procedure of implementing breeding program and fills in the blanks of the animal and plant breeding system.

